# Cognitive Benefits of Exercise Interventions: An fMRI Activation Likelihood Estimation Meta-Analysis

**DOI:** 10.1101/2020.07.04.187401

**Authors:** Qian Yu, Fabian Herold, Benjamin Becker, Ben KluGah-Brown, Yanjie Zhang, Stephane Perrey, Nicola Veronese, Notger G. Müller, Liye Zou, Arthur F. Kramer

**Affiliations:** Exercise & Mental Health Laboratory, School of Psychology, Shenzhen University, 518060, China; (QY); (YZ); (LZ); Research Group Neuroprotection, German Center for Neurodegenerative Diseases (DZNE), Leipziger Str. 44, 39120 Magdeburg, Germany; (F.H.); (NM); The Clinical Hospital of Chengdu Brain Science Institute, MOE Key Laboratory for Neuroinformation, University of Electronic Science and Technology of China, Chengdu 610054, Sichuan, China; (BB); (BK); Health and Exercise Science Laboratory, Institute of Sports Science, Seoul National University, Seoul, Republic of Korea; (YJZ); EuroMov, University of Montpellier, Montpellier, France; (SP); Primary Care Department, Azienda ULSS 3 (Unità Locale Socio Sanitaria) “Serenissima”, Dolo-Mirano District, Venice, Italy; (NV); Center for Cognitive and Brain Health, Department of Psychology, Northeastern University, Boston, MA 02115, USA. Email: (KA); Beckman Institute, University of Illinois at Urbana-Champaign, Champaign, IL 61801, USA

**Keywords:** Exercise, cognition, brain health, network

## Abstract

Despite a growing number of functional MRI studies reporting exercise-induced changes during cognitive processing, a systematic determination of the underlying neurobiological pathways is currently lacking. To this end, our neuroimaging meta-analysis included 20 studies and investigated the influence of exercise on cognition-related functional brain activation. The overall meta-analysis encompassing all experiments revealed exercise-induced changes in the left parietal lobe during cognitive processing. Subgroup analysis further revealed that in the younger-age group (<35 years old) exercise induced more widespread changes in the right hemisphere whereas in the older-age group (≥35 years old) exercise-induced changes were restricted to the left parietal lobe. Furthermore, subgroup analysis for exercise intervention duration, showed that shorter exercise interventions induced changes in regions connected with frontoparietal and default mode networks whereas regions exhibiting effects of longer interventions connected with frontoparietal and dorsal attention networks. Our findings suggest that physical exercise training leads to changes in functional activation patterns primarily located in precuneus and associated with frontoparietal, dorsal attention and default mode networks. Furthermore, exercise-induced changes in functional brain activation varied as a function of age and exercise intervention duration.

## 1. Introduction

Cognition covers a wide range of mental abilities that allow us to perceive, process and store information which, in turn, enable us to successfully interact with our environment ^1,2^. Consequently, cognitive performance is crucial to determine several aspects of successful everyday life functioning as well as health ^3,4^. Notably, there is increasing evidence in the literature showing that specific cognitive abilities such as processing speed and episodic memory gradually decline with aging ^4–7^, and poor cognitive performance leads to impairments in several aspects of everyday life, including walking, financial management, and driving ^8–10^ and has been associated with a strongly increased risk for neurological diseases such as dementia ^11,12^ and higher mortality risk ^13^. It is thus of general importance to preserve cognitive functions across the lifespan and particularly in aging populations.

Current approaches to preserve cognitive functions in the aging population focus on lifestyle changes and emphasize the role of regular physical activity and exercise ^14–17^. An increase in physical activity level is usually achieved through regular physical exercise (also referred to as physical training). Indeed, accumulating evidence indicates that both acute bout of physical exercise ^18–20^ and chronic exercise intervention ^19,21^ can influence cognitive performance positively. However, the underlying neurobiological processes which lead to an increase in cognitive performance after physical interventions are not fully understood.

According to Stillman ^22^, physical interventions induce changes on different levels of analysis which, in turn, promote the improvement of cognitive performance. In particular, physical exercise and physical training lead to changes on (i) molecular and cellular levels (e.g., brain-derived neurotrophic factors), (ii) structural and/or functional levels (e.g., hippocampus volume and hippocampal activity), and (iii) socioemotional level (e.g., sleep quality, well-being, self-efficacy) ^22^. Currently, there are systematic reviews and meta-analysis available which summarize the beneficial effects of acute physical exercise (a single bout of exercise) on changes: (i) molecular and cellular level (i.e., brain derived neurotrophic factor) ^23–26^, (ii) structural level ^27,28^ and (iii) socioemotional level (i.e., sleep) ^29,30^. In contrast the effects of exercise on functional brain changes that accompany acute physical exercise and chronic exercise intervention are currently less well understood. In this context, previous qualitative reviews have summarized the effects of acute ^31^ and chronic ^32,33^ physical exercise on functional brain changes, but did not perform a systematic quantitative meta-analytical analyses. Moreover, given that some theories of cognitive aging emphasize the importance of compensatory brain activation patterns in distinct functional neural networks ^34–37^, a deeper understanding of physical exercise-induced functional brain activation changes can help us to better tailor physical exercise interventions to individuals. Hence, this meta-analysis addresses this gap in the literature and investigates the influence of physical exercise interventions on cognition-related changes of functional brain activation.

## 2. Methods

This study followed the recommendations outlined in the Preferred Reporting Items for Systematic Review and Meta-Analysis (PRISMA) guidelines and has been registered on OSF Registries (Registration DOI: 10.17605/OSF.IO/674HF).

### 2.1 Data sources

The literature search was conducted at April 19^th^, 2020, through four electronic databases (Pubmed, Web of Science, PsychInfo, and Embase). Search terms in title and abstract were combined as follow:[movement OR “sport” OR “physical education” OR “physical activity” OR “aerobic exercise” OR “aerobic training” OR “interval training” OR “walking” OR “stretching*” OR “coordinative exercise” OR “coordinative training” OR “plyometric exercise” OR “plyometric training” OR “resistance exercise” OR “resistance training” OR “strength exercise” OR “strength training” OR “cardiopulmonary intervention” OR “cardiorespiratory training” OR “fitness training” OR “motor skill training” OR run OR cycle OR dance OR Tai Chi OR Yoga OR treadmill OR agility OR endurance OR “musculoskeletal intervention” OR “functional training” OR “physical therapy” OR “physiotherapy” OR exergam* OR “active gam*” OR “active play*” OR “interactive video” OR “virtual reality*” OR “motion gam*” OR Kinect OR wii*] AND [“cognition” OR “cognitive” OR “neuropsychological function” OR “executive function” OR “executive control” OR “central executive” OR “inhibitory control” OR “working memory” OR “task-switching” OR “planning” OR “attention” OR “information processing” OR “processing speed” OR “memory” OR “free recall” OR mental OR “emotion regulation” OR “cognitive regulation” OR “stress regulation” OR “psychological” OR “affective” brain OR “functional plasticity” OR “response time” OR “reaction time” OR accuracy OR error OR inhibition OR visual OR spatial OR visuospatial OR language OR oddball OR “problem solving” OR Flanker OR Stroop OR Sternberg] AND [fMRI OR MRI OR “MR imaging” OR “magnetic resonance imaging” OR “functional MRI” OR “functional magnetic resonance imaging” OR “PET” OR “positron emission tomography” OR “SPECT” OR “single-photon emission computed tomography”]. Furthermore, reference lists of included articles were manually searched for relevant articles that were captured through the database searches.

### 2.2 Inclusion criteria and study selection

The screening for relevant studies was conducted adhering to the PICOS-principles which stands for participants (P), intervention (I), comparisons (C), outcomes (O), and study design (S) ^38,39^. We included peer-review journal article published in English when they met the following inclusion criteria: (P) no restrictions were applied and we included all age groups regardless of pathologies; (I) only studies performing physical exercise/physical training were considered as eligible; (C) only studies with a pre/post intervention experimental designs and at least one group assigned to physical exercise/physical training intervention were included; (O) the relevant studies needed to assess cognition-related brain activation patterns via fMRI, PET or SPECT (task-based imaging studies) and needed to report retrievable data in standard Talairach or Montreal Neurologic Institute (MNI) coordinates. Based on the above-mentioned inclusion criteria, two independent researchers (QY and LZ) first screened article titles and abstracts to identify eligible articles. Afterwards and as recommended, a more detailed screening using the full-text of the article was conducted to ensure that all inclusion criteria were met.

### 2.3 Data extraction

The extraction of the relevant data was performed by two independent reviewers (QY and LZ) and the following information were extracted: (i) name of the lead author, (ii) imaging modality, (iii) population characteristics (e.g., health status and age), (iv) intervention characteristics (e.g., the number of participants, type of physical exercise, exercise duration, training frequency, training duration, and control condition), (v) cognitive task paradigms employed to assess the effects of the intervention, and (vi) functional brain activation results (e.g., the number of Foci). Risk of bias was independently assessed (by QY and LY) using the PEDro scale including 11 items ^40^.

### 2.4 Activation Likelihood Estimation (ALE)

In this study, we performed ALE analyses with GingerALE v3.0.2 (http://www.brainmap.org/ale/) ^41^ in MNI space, and cluster-based family-wise error (FWE) was used with a threshold of p < 0.05 (permuted 1000 times) ^41^. The p-value accounts for the proportion of the random spatial relation between the various experiments under the null distribution. Coordinates reported in Talairach space in the original studies were initially transformed to the MNI space using the Lancaster transform icbm2tal software procedure as implemented in the Convert Foci tool of GingerALE ^42^. The ALE maps were imported into Mango Version 4.1 (http://ric.uthscsa.edu/mango/mango.html) software and overlaid on an anatomical template in MNI space for visualization and comparison.

### 2.5 Anatomical connectivity, functional connectivity and functional characterization of the identified brain regions

In this study, if a meta-analytic identified cluster included more than one peak, we considered the area where the peak was centered at as sub-region (radius = 3mm). Anatomical and functional connection patterns of each sub-region were further determined by the Brainnetome Atlas and visualized by the Brainnetome Atlas Viewer (V1.0) ^43,44^. The functional characterizations of each sub-region are illustrated through probabilistic maps reflecting the behavioral domain and paradigm class according to meta data labels of the BrainMap database (http://www.brainmap.org/taxonomy). Overlapping behavioral domain(s) or paradigm(s) across sub-regions were subsequently selected ^43,44^. Brainnetome Atlas offers a fine-grained and cross-validated atlas providing structural information of more than 200 sub-regions. Brainnetome Atlas also maps the brain structure and function to mental processes by reference to the BrainMap database. Thus, Brainnetome Atlas provides an effective way for researchers to explore the complex relationship between anatomy, connectivity and function.

We also used the DPABI, a surface-based fMRI data analysis toolbox, to explore associations between regions/ sub-regions identified in both overall and subgroup analyses and the 7 networks proposed by Yeo et al ^45,46^. DPABI yoked between images were obtained from ALE analysis and DPABI template.

### 2.6 Subgroup Analysis

Given that exercise-induced changes on the brain functional and behavioral level might be influenced by both individual characteristics (e.g., health status, age and regular level of physical activity) and intervention characteristics (e.g., training duration), the included studies were categorized into the following subgroups (as recommended by previous reviews ^47,48^): (i) health status (healthy *vs*. patients) – the patient groups included studies conducted in individuals with social anxiety disorder, major depressive disorder, cognitive impairment, fibromyalgia, Parkinson’s disease, and bipolar disorder; (ii) age (“younger-age group: < 35 years old” *vs* “older-age group: ≥ 35 years old”). The cut-off of 35 years was employed to achieve a relatively balanced number of studies for each age-related subgroup analysis and was additionally based on evidence that cognitive function typically peaks around 35 years ^49^; (iii) training duration (“shorter-term: < 12 weeks” *vs* “longer-term: ≥ 12 weeks”). Based on previous meta-analysis, we used a 12-week training duration as the cut-off point for sub-group analysis ^48^; (iv) Although physical activity is characterized by three important components (intensity, frequency, and duration), it is reported that total amount of physical activity (i.e., minutes of exercise per week) is the most important factor for achieving health benefits ^48^. Hence, weekly total minutes of exercise were used to categorize studies into two groups, including physically inactive group (PIG) and physically active group (PAG). Specifically, we followed the criteria from the Physical Activity Guidelines for Americans (2nd Edition) to determine the group of each experiment-based study with the age-specific cut-off values: (a) 180 minutes per week in children and adolescents aged 6 to 17 years (PIG: < 180 minutes per week; PAG: ≥ 180 minutes per week); (b) 150 minutes (moderate-intensity aerobic exercise) per week in adults (PIG: < 150 minutes per week; PAG: ≥ 150 minutes per week). Notably, individuals aged over 65 years who kept on exercising were considered as physically active, as suggested by the aforementioned guideline ^48^.

## 3. Results

### 3.1 Study Selection

The systematic literature search returned a total of 42,302 records and 7,870 duplicates were removed (see Figure 1). The remaining 34,432 records were initially screened by examining the article titles and abstracts, with 34,195 records being excluded due to their failure to meet the pre-determined inclusion criteria (e.g., no original research or case reports, non-relevant outcomes). Full-text assessment of 237 was further conducted by two independent authors (QY and LZ), which resulted in 20 eligible studies; as shown in Figure 1,217 records were excluded according to our selection criteria (review and conference abstract = 24, no pre-to-post imaging assessment = 10, irrelevant outcomes = 168, no coordinates of whole-brain analysis = 3, non-cognitive task = 2; non-exercise intervention = 10).

**Figure 1.**
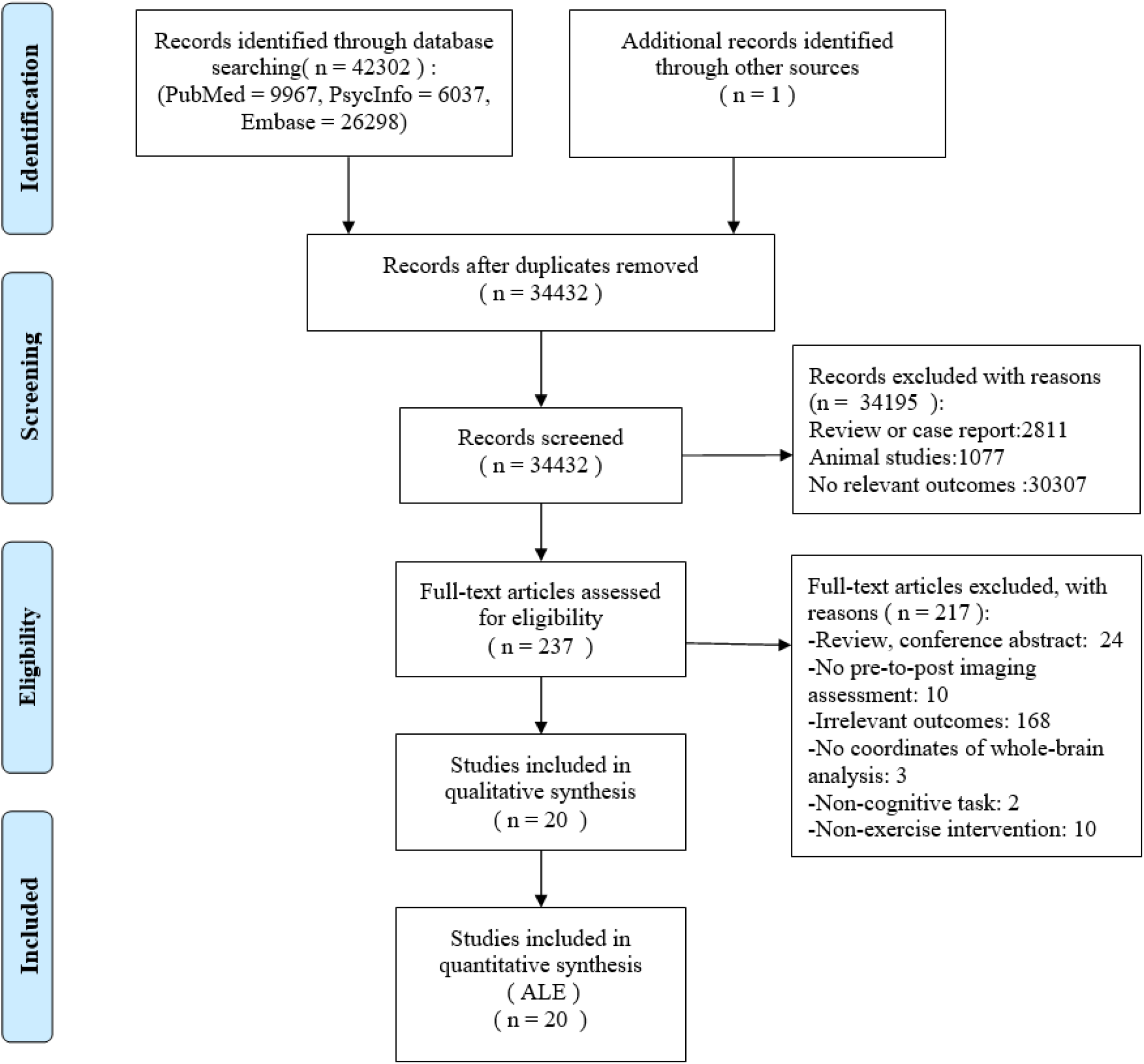
Flow Chart of Literature Searching, Screening and Assessment

### 3.2 Characteristics of Included Studies

There are 260 foci of activation within 20 studies including a total 745 participants with mean age of 47.41 (SD = 22.04): 283 patients (mean age = 44.01, SD = 20.28) and 463 healthy people (mean age = 49.49, SD = 22.83). Study characteristics are detailed in Table 1.

**Table 1.** Information of included studies.

| No. | Study | Mode | Population | Age | Intervention of Experimental Group | Duration | Task/ Stimuli | Control | Foci |
| --- | --- | --- | --- | --- | --- | --- | --- | --- | --- |
| 1 | Goldin et al., 2012 (a)<sup>50</sup> | fMRI | Patients with social anxiety disorder | 32.88±7.97 | Aerobic exercise (n=18) | 2.5 hours/week * 8 weeks (weekly at least two individual AE sessions and one group AE session, nearly 2.5 hours) | Self-referential encoding task (self-referent) | Mindfulness-based stress reduction (n=24) | 15 |
| 2 | Goldin et al., 2012 (b)<sup>51</sup> | fMRI | Patients with social anxiety disorder | 32.88±7.97 | Aerobic exercise (n=19) | 2.5 hours/week * 8 weeks (weekly at least two individual AE sessions and one group AE session, nearly 2.5 hours) | Cognitive reappraisal task (negative self-belief) | Mindfulness-based stress reduction (n=23) | 6 |
| 3 | Gourgouvelis et al., 2017<sup>52</sup> | fMRI | Patients with major depressive disorder | 37.25±8.00 | Aerobic exercise for patients(n=8) | 60min/session * 1 session/week * 8 weeks | Associative memory task (memory) | Aerobic exercise for healthy people (n=8) | 10 |
| 4 | Schmitt et al., 2019<sup>53</sup> | fMRI | Athletes | 27.20±4.20 | low- vs high-intensity exercise (n=21) | 40 min for low-intensity exercise<br>40 min for high-intensity exercise | fMRI-based face-matching paradigm (fear/ happiness) | / | 4 |
| 5 | Boa Sorte Silva et al., 2020<sup>54</sup> | fMRI | Older adults | 67.50±7.30 | multiple-modality exercise+ mind-motor training (n=63) | 60 min/day * 3 days/week * 24 weeks | Computer-based memory tasks (working & episodic memory) | multiple-modality exercise+ others (n=64) | 38 |
| 6 | Hsu et al., 2016<sup>55</sup> | fMRI | Older adults with vascular cognitive impairment | 72.60±8.90 | Aerobic exercise (n=10) | 60 min/session * 3 sessions/week * 24 weeks | Eriksen flanker task (executive function) | Usual care (n=11) | 16 |
| 7 | Li et al., 2019 <sup>56</sup> | fMRI | Female college students | 25.50±0.67 | Acute aerobic exercise for high-fitness students (n=12) | 30 min | N-back task (executive function) | Acute aerobic exercise for low-fitness students (n=12) | 5 |
| 8 | Martinsen et al., 2017 <sup>57</sup> | fMRI | Patients with fibromyalgia | 25-64y | Physical exercise for patients (n=19) | 60 min/session * 2 sessions/week * 15 weeks | Stroop colour word test (executive function) | Physical exercise for healthy people (n=20) | 3 |
| 9 | Nishiguchi et al., 2015 <sup>58</sup> | fMRI | Older adults | 73.00±4.80 | Dual-task group exercise (n=24) | 90 min/session * 1 session/week * 12 weeks | N-back task (working memory) | Usual care (n=24) | 7 |
| 10 | Smith et al., 2013 <sup>59</sup> | fMRI | Patients with mild cognitive impairment | 78.70±7.50 | Moderate intensity treadmill walking for mild cognitive impairment patients (n=17) | 30 min/session * 44 sessions, for 12 weeks | Famous name discrimination task (semantic memory) | Moderate intensity treadmill walking for healthy people (n=18) | 26 |
| 11 | Krafft et al., 2014 <sup>60</sup> | fMRI | Overweight children | 9.70±0.80 | Aerobic exercise (n=24) | 40 min/session * 138 sessions, for 8 months | Antisaccade task<br>Flanker task (cognitive control) | Attention control (n=19) | 10 |
| 12 | Duchesne et al., 2016 <sup>61</sup> | fMRI | Patients with Parkinson's disease | 59.00±7.11 | Aerobic exercise for Parkinson's disease patients (n=19) | 20 min/session (+ 5 min/week up to 40 min) * 3 sessions/week * 12 weeks | Serial reaction time task (procedural memory) | Aerobic exercise for healthy people (n=19) | 8 |
| 13 | Chen et al., 2016 <sup>62</sup> | fMRI | Preadolescent children | 10y | Acute aerobic exercise (n=9) | 35 min | N-back task (working memory) | / | 5 |
| 14 | Pensel et al., 2018 <sup>63</sup> | fMRI | Middle-aged sedentary males | 49.00±5.32 | Exercise training (n=23) | 90 min/session * 3 sessions/week * 24 weeks | Flanker paradigm (Executive control processing) | Sedentary lifestyle (n=14) | 32 |
| 15 | Liu-Ambrose et al., 2012 <sup>64</sup> | fMRI | Community-dwelling senior women | 69.3±3.2y | Once-weekly resistance training (n1=20)<br>Twice-weekly resistance training (n2=15) | 1 month (once-weekly/<br>twice weekly) | Modified Eriksen flanker task (selective attention and conflict resolution) | Twice-weekly balance and tone training (n =17) | 12 |
| 16 | Wagner et al., 2017 <sup>65</sup> | fMRI | Young healthy male | 25.0±3.3y | Short-term intense aerobic exercise (n=17) | 60min/session * 3 sessions/week * 6 weeks | Paired-associated learning task (Memory) | No additional specific training (n=17) | 36 |
| 17 | Wu et al., 2018 <sup>66</sup> | fMRI | Healthy people | 64.9±2.8y | Yang-style Tai Chi Chuan (n=16) | 60min/week * 12 weeks | Numerical stroop paradigm (Executive function: Task-switching performance) | Telephone consultation (n=10) | 5 |
| 18 | Wriessnegger et al., 2014 <sup>67</sup> | fMRI | Healthy people | 28.4±4.3y | Playing soccer and tennis via "Kinect" (n=23) | 20min | Imagery task (Motor imagery) | / | 8 |
| 19 | Metcalfe et al., 2016 <sup>68</sup> | fMRI | Adolescents with bipolar disorder | 16.8±1.4y | Acute aerobic exercise: cycling (n=30) | 27min | Sustained attention to response task (Attention) | Acute aerobic exercise for healthy people (n=20) | 8 |
| 20 | Baeck et al., 2012 <sup>69</sup> | fMRI | Healthy people | 27.2±7.9 | Shooting training (n=18) | 90min/session * 3 sessions/week * 20 weeks | Motor imagery paradigm (Motor imagery) | / | 6 |

### 3.3 Overall Analysis: Exercise-Induced Brain Activation Associated with Cognition

Twenty studies that investigated the effects of physical exercise on cognition-associated functional brain activation were included in this meta-analysis ^50–69^. Overall, 260 foci from all 26 experiments converged onto a 2200mm^3^ cluster centered at (−26.2, −57.9, 45.3) with 4 peaks (Figure 2: (a)). All 4 peaks were located at the parietal lobe of left cerebrum. More specifically, the cluster encompassed regions in the precuneus (69.7%), inferior parietal lobule (27.3%), and superior parietal lobule (3%) and covered regions no Brodmann areas 7, 19, 39, and 40.

Examining the direction of the effects in terms of increased activation, 184 foci from 17 experiments were included and converged on a 1616 mm^3^ cluster centered at (38.1, −21.9, 4.9) with 3 peaks (Figure 2: (b)) covering sub-lobar and temporal lobe regions in the right brain hemisphere. The cluster was primarily located in the lentiform nucleus (71.2%) and spread in the adjacent claustrum (17.8%), superior temporal gyrus (6.8%), as well as the insula (4.1%). Accordingly, the activated cluster comprises the Brodmann areas 13, 22 and 41 as well as the putamen. With respect to decreased activation reported in 9 experiments, no region of convergently decreased activation was identified.

**Figure 2.**
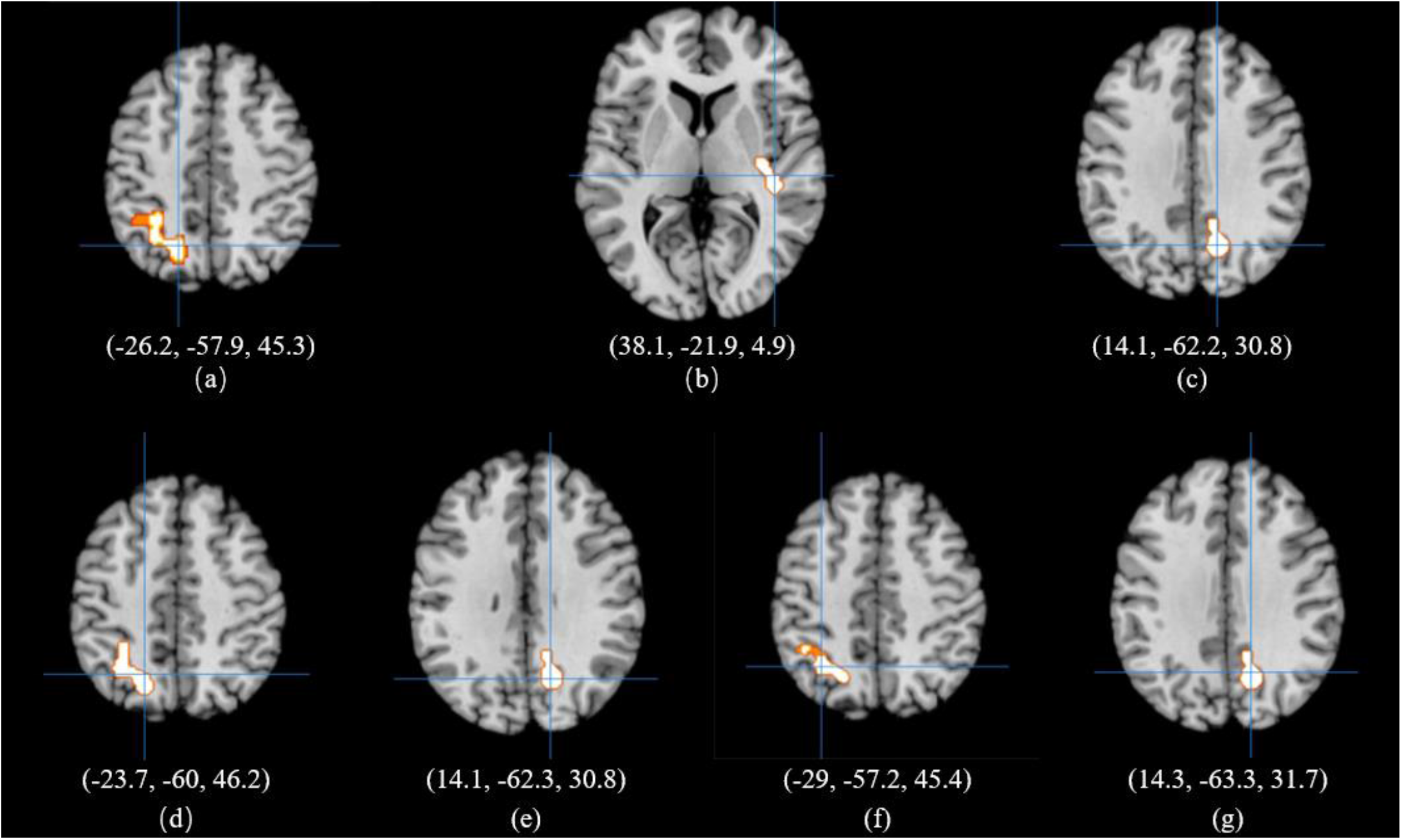
Activated Clusters in Overall Analysis and Subgroup Analyses Notes: (a) activation for overall analysis; (b) increased activation; (c) activation for <35-year-old subgroup; (d) activation for ≥35-year-old subgroup; (e) activation for <12-week subgroup; (f) activation for ≥12-week subgroup; (g) activation for physically inactive subgroup.

### 3.4 Subgroup Analysis for Health Status

For the healthy population ^53,54,56,58,60,62–67,69^, 168 foci from 14 experiments were included for the subgroup analysis; and for the patient population ^50–52,55,57,59,61,68^, 92 foci from 12 experiments were selected. There was no activated cluster for either analysis.

### 3.5 Subgroup Analysis for Age

In the younger-age group ^50,51,53,56,60,62,65,67–69^, 103 activation foci from 15 experiments were included and converged on 2112mm^3^ cluster centered at (14.1, −62.2, 30.8) with 3 peaks (Figure 2: (c)) located at the occipital, parietal and limbic lobe. The identified cluster primarily encompassed the precuneus (78.5%), spreading into the adjacent cuneus (10.3%), posterior cingulate (6.5%), and cingulate gyrus (4.7%). The cluster encompassed regions located in Brodmann areas 7 and 31.

In the older-age group ^52,54,55,57–59,61,63,64,66^, 157 foci from 11 experiments were included and converged on a 2360 mm^3^ cluster spanning from (−36, −72, 40) to (−10, −44, 56), with the center at (−23.7, −60, 46.2) (Figure 2: (d)). The 4 peaks of this cluster were located in the parietal lobe of the left hemisphere. The identified cluster was mainly located in the precuneus (82.2%), and additionally encompassed the inferior parietal lobule (6.7%), superior parietal lobule (6.7%), and angular gyrus (4.4%). In addition, the cluster included regions located in Brodmann areas 7, 19 and 39.

### 3.6 Subgroup Analysis for Intervention Duration

For the shorter-term duration interventions ^50–53,56,62,64,65,67,68^, 109 foci from 14 experiments converged on a 2008mm^3^ cluster centered at (14.1, −62.3, 30.8) with 3 peaks in right hemisphere (Figure 2: (e)). Among these peaks, 1 peak with the maximum value was located in the occipital lobe and 2 peaks were situated in the limbic lobe. The cluster was primarily located in the precuneus (80.2%), spreading into the cuneus (10.9%), posterior cingulate (5%), and cingulate gyrus (4%). At the brain region level, Brodmann areas 31 and 7 contributed to 73.3% and 25.7% to this cluster, respectively. For the longer-term duration interventions ^54,55,57–61,63,66,69^, 151 foci from 12 experiments converged on a 1968mm^3^ cluster centered at (−29, −57.2, 45.4) with 4 peaks in the parietal lobe of left cerebrum (Figure 2: (f)). The cluster encompassed the precuneus (45.5%), as well as the inferior parietal lobule (45.5%), angular gyrus (6.1%), and supramarginal gyrus (3%) corresponding to Brodmann areas 7, 19, 39 and 40.

### 3.7 Subgroup Analysis for Total Amount of Physical Activity

In the PIG ^52,53,55,57,58,60–62,64,66–68^, 85 foci from 16 experiments converged on a 1792mm^3^ cluster centered at (14.3, −63.3, 31.7) with 2 peaks (Figure 2: (g)). Peaks 1 and 2 were located in the right occipital lobe and in the right limbic lobe, respectively. The cluster covered the occipital lobe, parietal lobe and limbic lobes. The cluster primarily included the precuneus (83.8%), and cuneus (33.3%) with additional engagement of the cingulate gyrus (5.1%), which correspond to Brodmann area 31 (72.7%) and Brodmann area 7 (27.3%). In the PAG ^50,51,54,55,59,63,65,69^, 175 foci from 10 experiments did not converge on a robust cluster.

### 3.8 Anatomical Connectivity, Functional Connectivity and Functional Characterizations of Activated Brain Regions

In this study, each activated cluster includes more than one peak with each one that can generate a sub-region (radius = 3mm). Anatomical and functional connectivity of activated sub-regions (4 peaks) in the overall analysis and subgroup analyses are shown in Figure 3. Functional characterizations of activated sub-regions in both overall analysis and subgroup analyses are summarized in Table 2, and revealed that the identified sub-regions in the overall analysis exhibited a strong positive coupling with the entire frontoparietal control network and were functionally characterized by engagement in core cognitive domains, including attention, executive functions and working memory. For the functional characterizations, sub-regions were connected with spatial cognition in the overall analysis. In the subgroup analyses, activated sub-regions were associated with explicit memory in the younger-age group, shorter-term group, and PIG, whereas activated sub-regions were associated with working memory in the older-age group and longer-term intervention group. In addition, in both overall analysis and subgroup analysis (older-age group), activated sub-regions were associated with paradigms measuring mental rotation.

**Figure 3.**
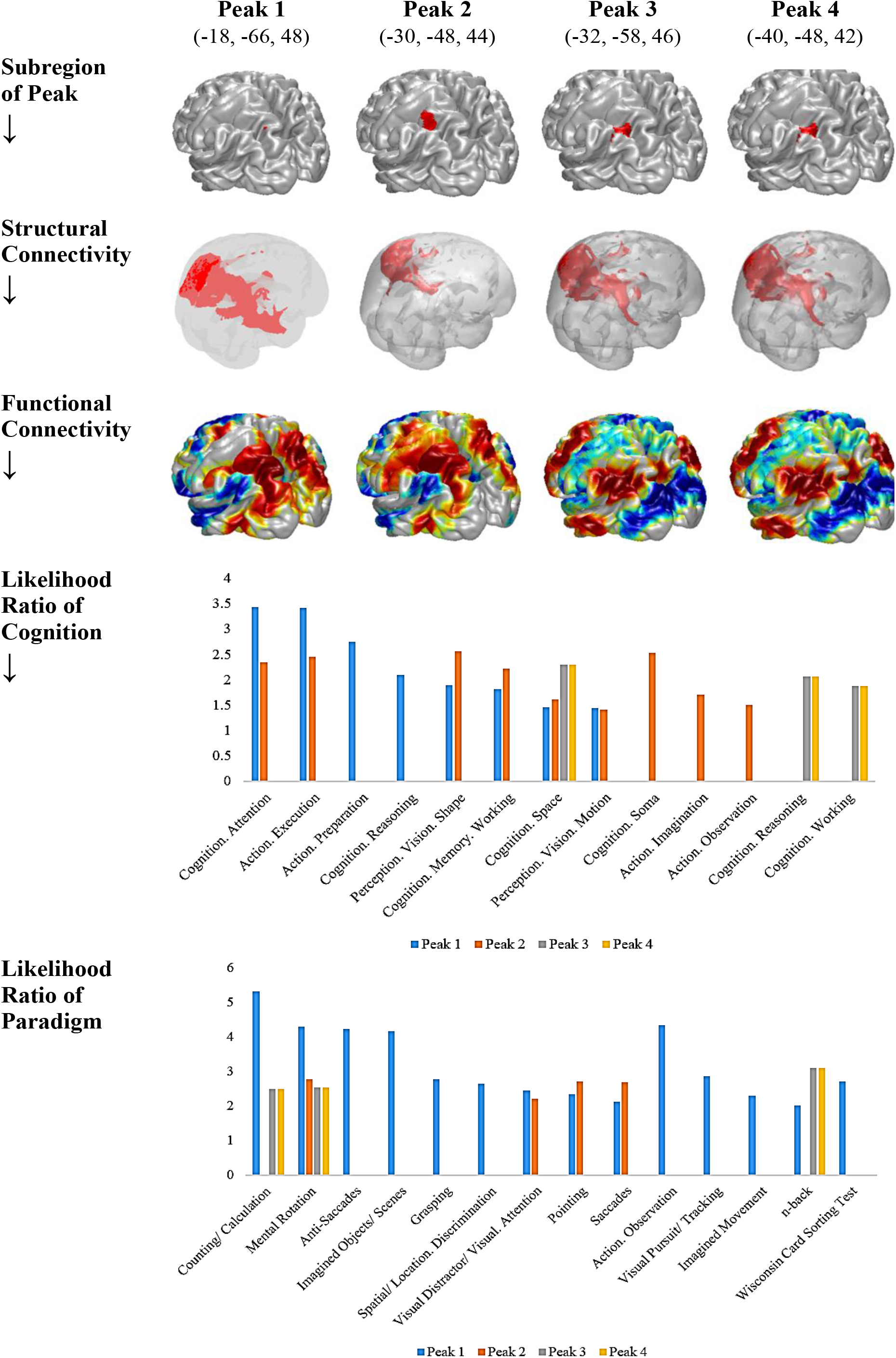
Anatomical Connectivity, Functional Connectivity and Functional Characterizations of Activated Brain sub-regions in the Overall Analysis

**Table 2 (a).** Characterizations of activated brain regions in overall analysis.

|  | Coordinate |  |  | Location | Likelihood Ratio of Behavioral Domain |  | Likelihood Ratio of Paradigm |  |
| --- | --- | --- | --- | --- | --- | --- | --- | --- |
|  | x | y | z |  |  |  |  |  |
| **Peak 1** | -18 | -66 | 48 | Left Cerebrum. Parietal Lobe. Precuneus. Gray Matter. Brodmann area 7 | Cognition. Attention | 3.44 | Counting/ Calculation | 5.31 |
|  |  |  |  |  | Action. Execution | 3.41 | Mental Rotation© | 4.28 |
|  |  |  |  |  | Action. Preparation | 2.75 | Anti-Saccades | 4.23 |
|  |  |  |  |  | Cognition. Reasoning | 2.10 | Imagined Objects/ Scenes | 4.15 |
|  |  |  |  |  | Perception. Vision. Shape | 1.90 | Grasping | 2.77 |
|  |  |  |  |  | Cognition. Memory. Working | 1.82 | Spatial/ Location. Discrimination | 2.65 |
|  |  |  |  |  | Cognition. Space ® | 1.46 | Visual Distractor/ Visual. Attention | 2.45 |
|  |  |  |  |  | Perception. Vision. Motion | 1.45 | Pointing | 2.33 |
|  |  |  |  |  |  |  | Saccades | 2.11 |
| **Peak 2** | -30 | -48 | 44 | Left Cerebrum. Parietal Lobe. Superior Parietal Lobule. Gray Matter. Brodmann area 7 | Perception. Vision. Shape | 2.56 | Action. Observation | 4.34 |
|  |  |  |  |  | Cognition. Soma | 2.53 | Visual Pursuit/ Tracking | 2.85 |
|  |  |  |  |  | Action. Execution | 2.45 | Mental Rotation© | 2.77 |
|  |  |  |  |  | Cognition. Attention | 2.35 | Pointing | 2.70 |
|  |  |  |  |  | Cognition. Memory. Working | 2.22 | Saccades | 2.69 |
|  |  |  |  |  | Action. Imagination | 1.71 | Imagined Movement | 2.29 |
|  |  |  |  |  | Cognition. Space ® | 1.61 | Visual Distractor/ Visual. Attention | 2.20 |
|  |  |  |  |  | Action. Observation | 1.51 | n-back | 2.01 |
|  |  |  |  |  | Perception. Vision. Motion | 1.41 |  |  |
| **Peak 3** | -32 | -58 | 46 | Left Cerebrum. Parietal Lobe. Inferior Parietal Lobule. Gray Matter. Brodmann area 39 | Cognition. Space ® | 2.29 | n-back | 3.1 |
|  |  |  |  |  | Cognition. Reasoning | 2.06 | Wisconsin Card Sorting Test | 2.71 |
|  |  |  |  |  | Cognition. Working | 1.88 | Mental Rotation© | 2.53 |
|  |  |  |  |  |  |  | Counting/ Calculation | 2.49 |
| **Peak 4** | -40 | -48 | 42 | Left Cerebrum. Parietal Lobe. Inferior Parietal Lobule. Gray Matter. Brodmann area 40 | Cognition. Space ® | 2.29 | n-back | 3.1 |
|  |  |  |  |  | Cognition. Reasoning | 2.06 | Wisconsin Card Sorting Test | 2.71 |
|  |  |  |  |  | Cognition. Working | 1.88 | Mental Rotation© | 2.53 |
|  |  |  |  |  |  |  | Counting/ Calculation | 2.49 |

Note: ® and © indicate overlapping behavioral domain(s) or paradigm(s) across sub-regions, respectively.

**Table 2 (b).** Characterizations of activated brain regions in subgroup analyses.

|  | Coordinate |  |  | Location | Likelihood Ratio of Behavioral Domain |  | Likelihood Ratio of Paradigm |  |
| --- | --- | --- | --- | --- | --- | --- | --- | --- |
|  | x | y | z |  |  |  |  |  |
| < 35 Years Old Subgroup |  |  |  |  |  |  |  |  |
| Peak 1 | 14 | -64 | 32 | Right Cerebrum. Occipital Lobe. Precuneus. Gray Matter. Brodmann area 31 | Perception. Vision. Motion | 2.06 | Saccades | 2.51 |
|  |  |  |  |  | Cognition. Memory. Explicit® | 1.94 |  |  |
| Peak 2 | 12 | -52 | 32 | Right Cerebrum. Limbic Lobe. Cingulate Gyrus. Gray Matter. Brodmann area 31 | Cognition. Social Cognition | 3.11 | Cued Explicit. Recognition | 5.88 |
|  |  |  |  |  | Cognition. Memory. Explicit® | 2.45 | Episodic Recall | 3.48 |
|  |  |  |  |  |  |  | Theory of Mind Task | 2.46 |
| Peak 3 | 12 | -54 | 22 | Right Cerebrum. Limbic Lobe. Posterior Cingulate. Gray Matter. Brodmann area 23 | Perception. Vision. Motion | 2.06 | Saccades | 2.51 |
|  |  |  |  |  | Cognition. Memory. Explicit® | 1.94 |  |  |
| ≥35 Years Old Subgroup |  |  |  |  |  |  |  |  |
| Peak 1 | -18 | -68 | 48 | Left Cerebrum. Parietal Lobe. Precuneus. Gray Matter. Brodmann area 7 | Cognition. Attention | 3.44 | Counting/ Calculation | 5.31 |
|  |  |  |  |  | Action. Execution | 3.41 | Mental Rotation | 4.28 |
|  |  |  |  |  | Action. Preparation | 2.75 | Anti-Saccades | 4.23 |
|  |  |  |  |  | Cognition. Reasoning | 2.10 | Imagined Objects/ Scenes | 4.15 |
|  |  |  |  |  | Perception. Vision. Shape | 1.90 | Grasping | 2.77 |
|  |  |  |  |  | Cognition. Memory. Working® | 1.82 | Spatial/ Location. Discrimination | 2.65 |
|  |  |  |  |  | Cognition. Space® | 1.46 | Visual Distractor/ Visual. Attention | 2.45 |
|  |  |  |  |  | Perception. Vision. Motion | 1.45 | Pointing | 2.33 |
|  |  |  |  |  |  |  | Saccades | 2.11 |
| Peak 2 | -30 | -48 | 44 | Left Cerebrum. Parietal Lobe. Superior Parietal Lobule. Gray Matter. Brodmann area 7 | Perception. Vision. Shape | 2.56 | Action. Observation | 4.34 |
|  |  |  |  |  | Cognition. Soma | 2.53 | Visual Pursuit/ Tracking | 2.85 |
|  |  |  |  |  | Action. Execution | 2.45 | Mental Rotation | 2.77 |
|  |  |  |  |  | Cognition. Attention | 2.35 | Pointing | 2.70 |
|  |  |  |  |  | Cognition. Memory. Working® | 2.22 | Saccades | 2.69 |
|  |  |  |  |  | Action. Imagination | 1.71 | Imagined Movement | 2.29 |
|  |  |  |  |  | Cognition. Space® | 1.61 | Visual Distractor/ Visual. Attention | 2.20 |
|  |  |  |  |  | Action. Observation | 1.51 | n-back | 2.01 |
|  |  |  |  |  | Perception. Vision. Motion | 1.41 |  |  |
| Peak 3 | -32 | -58 | 46 | Left Cerebrum. Parietal Lobe. Inferior Parietal Lobule. Gray Matter. Brodmann area 39 | Cognition. Space® | 2.29 | n-back | 3.10 |
|  |  |  |  |  | Cognition. Reasoning | 2.06 | Wisconsin Card Sorting Test | 2.71 |
|  |  |  |  |  | Cognition. Memory. Working® | 1.88 | Mental Rotation | 2.53 |
|  |  |  |  |  |  |  | Counting/ Calculation | 2.49 |
| Peak 4 | -12 | -64 | 54 | Left Cerebrum. Parietal Lobe. Precuneus. Gray Matter. Brodmann area 7 | / | / | / | / |
| < 12 Weeks Subgroup |  |  |  |  |  |  |  |  |
| Peak 1 | 14 | -64 | 32 | Right Cerebrum. Occipital Lobe. Precuneus. Gray Matter. Brodmann area 31 | Perception. Vision. Motion | 2.06 | Saccades | 2.51 |
|  |  |  |  |  | Cognition. Memory. Explicit® | 1.94 |  |  |
| Peak 2 | 12 | -52 | 32 | Right Cerebrum. Limbic Lobe. Cingulate Gyrus. Gray Matter. Brodmann area 31 | Cognition. Social Cognition | 3.11 | Cued Explicit. Recognition | 5.88 |
|  |  |  |  |  | Cognition. Memory. Explicit® | 2.45 | Episodic Recall | 3.48 |
|  |  |  |  |  |  |  | Theory of Mind Task | 2.46 |
| Peak 3 | 12 | -54 | 22 | Right Cerebrum. Limbic Lobe. Posterior Cingulate. Gray Matter. Brodmann area 23 | Perception. Vision. Motion | 2.06 | Saccades | 2.51 |
|  |  |  |  |  | Cognition. Memory. Explicit® | 1.94 |  |  |
| ≥12 Weeks Subgroup |  |  |  |  |  |  |  |  |
| Peak 1 | -32 | -58 | 46 | Left Cerebrum. Parietal Lobe. Inferior Parietal Lobule. Gray Matter. Brodmann area 39 | Cognition. Space® | 2.29 | n-back | 3.10 |
|  |  |  |  |  | Cognition. Reasoning® | 2.06 | Wisconsin Card Sorting Test | 2.71 |
|  |  |  |  |  | Cognition. Memory. Working® | 1.88 | Mental Rotation | 2.53 |
|  |  |  |  |  |  |  | Counting/ Calculation | 2.49 |
| Peak 2 | -40 | -48 | 42 | Left Cerebrum. Parietal Lobe. Inferior Parietal Lobule. Gray Matter. Brodmann area 40 | Cognition. Space® | 2.29 | n-back | 3.10 |
|  |  |  |  |  | Cognition. Reasoning® | 2.06 | Wisconsin Card Sorting Test | 2.71 |
|  |  |  |  |  | Cognition. Memory. Working® | 1.88 | Mental Rotation | 2.53 |
|  |  |  |  |  |  |  | Counting/ Calculation | 2.49 |
| Peak 3 | -20 | -64 | 46 | Left Cerebrum. Parietal Lobe. Precuneus. Gray Matter. Brodmann area 7 | Cognition. Reasoning® | 3.25 | Saccades | 3.54 |
|  |  |  |  |  | Cognition. Memory. Working® | 2.92 | Delayed Match To Sample | 3.37 |
|  |  |  |  |  | Cognition. Attention | 2.81 | Stroop Task | 2.73 |
|  |  |  |  |  | Perception. Vision. Motion | 2.37 | Mental Rotation | 2.68 |
|  |  |  |  |  | Cognition. Space® | 1.72 | Wisconsin Card Sorting Test | 2.45 |
|  |  |  |  |  | Perception. Vision. Shape | 1.68 | Task Switching | 2.44 |
| Peak 4 | -12 | -64 | 54 | Left Cerebrum. Parietal Lobe. Precuneus. Gray Matter. Brodmann area 7 | / | / | / | / |
| Physically Inactive Subgroup |  |  |  |  |  |  |  |  |
| Peak 1 | 14 | -64 | 32 |  | Perception. Vision. Motion | 2.06 | Saccades | 2.51 |
|  |  |  |  | Right Cerebrum. Occipital Lobe.<br>Precuneus. Gray Matter. Brodmann area 31 | Cognition. Memory. Explicit® | 1.94 |  |  |
| **Peak 2** | 12 | -52 | 32 | Right Cerebrum. Limbic Lobe. Cingulate<br>Gyrus. Gray Matter. Brodmann area 31 | Cognition. Social Cognition | 3.11 | Cue Explicit. Recognition | 5.88 |
|  |  |  |  |  | Cognition. Memory. Explicit® | 2.45 | Episodic Recall | 3.48 |
|  |  |  |  |  |  |  | Theory of Mind Task | 2.46 |

Note: ® and © indicate overlapping behavioral domain(s) or paradigm(s) across sub-regions, respectively.

In the overall analysis, the frontoparietal and dorsal attention networks were involved in exercise-induced changes in functional activation patterns. For subgroup analyses, associated networks varied across subgroups: (1) younger-age group (frontoparietal network) and older-age group (dorsal attention network); (2) shorter-term group (frontoparietal and default networks) and longer-term group (frontoparietal and dorsal attention networks); (3) PIG (frontoparietal and default networks).

### 3.9 Risk of Bias Assessment

Total score across included studies ranged from 3 to 7 (M = 4.80 and SD = 1.40) that correspond to poor to good quality. Notably, only two studies scored 7. Points in the majority of included studies were mainly deducted due to their study design such as lack of random allocation (n = 9), concealed allocation (n =14), assessor blinding (n = 18), intention-to-treat analysis (n = 18). Detailed information is displayed in Supplementary data.

## 4 Discussion

### 4.1 Exercise-Induced Brain Activation Associated with Cognition

The overall meta-analysis encompassing data from all original studies demonstrated robust exercise-induced changes in cognition related functional activation of the left parietal lobe, primarily covering the precuneus (69.7%) and spreading into inferior (27.3%) and superior (3%) parietal lobe. The precuneus plays a key role in a range of highly integrated mental processes, ranging from basic cognitive processes to regulatory control over performance under stress ^70,71^. The integrative function of the precuneus is further reflected in the functional connectivity profiles of these regions, which include separable interactions with networks engaged in sensorimotor and cognitive processes (e.g., executive function, episodic memory, visuospatial processing) ^72^. Sporns and Bullmore proposed that the precuneus plays a key role in in the frontoparietal network by interconnecting parietal and prefrontal regions (“small-world network” hub), which provides an explanation for the aforementioned activation during cognitive tasks ^73^. Moreover, the precuneus is a brain region which is commonly affected in individuals with mild cognitive impairment and early stages of Alzheimer’s disease ^74^. In light of this observation, our findings suggest that regular physical exercise can be a valuable approach to prevent cognitive decline ^15,16,75^ by enhancing the functional integration of the frontoparietal control network via effects on the precuneus.

The activated sub-regions were connected with dorsal attention network and frontoparietal network, and the functional characterizations revealed a strong engagement in core cognitive domains, including attention as well as executive functions. The frontoparietal network is a functional hub sharing connectivity with diverse brain networks and plays an essential role in modulating cognitive control ^76^. Moreover, the degree of frontoparietal network’s coupling with other brain networks (especially default mode network) is positively correlated with fluid intelligence (e.g., problem solving/executive function and visuospatial ability) and overall cognitive ability ^77,78^, which is consistent with our results in overall analysis for global cognition (all cognitive components were pooled together for the overall analysis). The dorsal attention network, centered around the intraparietal sulcus and frontal eye fields, participates in top-down control of attention as well as sensory-motor information integration, which support several higher level cognitive domains ^79^.

### 4.2 Moderators have impact on exercise-induced cognitive improvement

In this review, age and training duration considerably moderated exercise effects on brain activation that is associated with anatomical and functional networks. For instance, in the younger-age group, the identified sub-regions are associated with explicit memory. This result is strongly supported by a recent meta-analysis (including 25 experimental studies) demonstrating robust effects of exercise on episodic memory function ^80^. Furthermore, the recruited participants in 23 studies were reported with age range of 18 to 30 years old, which coincides with the younger-age group (less than 35 years old) in the current review. In the older-age group, identified sub-regions are related to executive functions, including working memory, spatial cognition and mental rotation. Such improvements induced by exercise have been well-documented in previous meta-analytical studies at the behavioral level ^21,81,82^.

With regard to the moderating effect of training duration on exercise-induced activation, shorter-term exercise interventions induced primarily changes in occipital, parietal and limbic regions and the peaks/sub-regions are generally linked to explicit memory, whereas the longer-term training protocols induced changes in activation in the parietal lobe. Of note, the shorter-term group included 14 experiments with 7 using acute physical exercise, which is partially supported by results from a previous meta-analysis which investigated the beneficial effects of acute exercise on episodic memory as a type of explicit memory ^83^. Meanwhile, researchers emphasized that the timing of acute exercise plays an important role in the interaction of exercise and memory; such that cognitive improvement was observed with acute exercise occurring before memory encoding, during early memory consolidation and during late memory consolidation. Positive results on explicit memory function may be attributed to improved pattern separation ^84,85^ and attenuated memory interference ^86^ following a relatively short intervention duration. For the longer-term group, the activated cluster was located in the parietal lobe and peak areas were associated with spatial processing, reasoning and working memory. For longer-term exercise, various positive effects on cognition have been reported at the molecular and cellular levels (e.g., brain-derived neurotrophic factor) ^87^, structural level (e.g., increased gray matter volume in frontal and hippocampal regions) ^88^, and behavioral level (e.g., improvement in executive function) ^89,90^.

Of note, the analyses separately examining healthy subjects and patient groups did not converge on a specific brain region that exhibited changes. However, when lowering the statistsical threshold (uncorrected) convergent activiation in the parietal lobe emerged for both, healthy subjects and patients (detailed information can be found in supplementary data), indicating that the lack of convergent activiation in the subgroups might be due to the reduced number of included experiments in each subgroup analysis. Besides, previous reviews have shown that both healthy people ^91^ and individuals with neurological, non-neurological, and psychiatric illnesses ^92,93^ can benefit from physical exercise on the behavioral level. For instance, a meta-analysis by Meenakshi et al, focusing on individuals with chronic disorders (Alzheimer’s disease, Huntington’s disease, multiple sclerosis, Parkinson's disease, schizophrenia, and unipolar depression), suggested that exercise interventions can improve several cognitive domains (attention, working memory and executive function) with small but significant effect sizes. In addition, an activated cluster was only observed in PIG, but not in PAG. Such results might be explained by physiological and psychological (cognitive) adaptation, which refers to that cellular stress and the resultant metabolic signals have reached relatively stable status ^94^. It is well known that fMRI measures brain activation by detecting blood-oxygen-level dependent response to cognitive tasks. If the maximum cognitive benefits are achieved with a total amount of 150 minutes or below, activation may be no longer present with greater amount of physical activity/exercise because of adaptation related to blood-oxygen metabolism ^95^.

### 4.5 Implications

This is the first study to systematically examine the relationship between functional brain activation and cognition as a function of exercise practice. Therefore, the present results can be used to guide future research on exercise effects on brain health and cognition. Firstly, in the subgroup analyses of age and training duration, activation was observed in different brain regions. Follow-up studies should explore the associations between hemispheric lateralization and moderators (age and duration). Secondly, only the improvement of spatial cognition was observed in the overall analysis without consideration of moderators, while the benefits on various cognitive components like explicit memory, working memory and reasoning were found in the subgroup analyses. It can be inferred that the cognitive benefits from exercise participation are specifically influenced by age and training duration, which is consistent with previous behavioral meta-analytical reviews ^96,97^. Thirdly, because of the limited number of studies available for analysis, we cannot determine the influences of sex, exercise type and disease type on cognitive improvement. Therefore, these moderators should be examined in future exercise-cognition studies using neuroimaging techniques. Fourthly, the small number of included experiments has limited researchers to focus on two age groups only for subgroup analysis. Furthermore, original imaging studies on this topic that focused on children and adolescent, middle-aged people, young-old (55-65 years old) group, oldest-old population are still in its infancy, which requires further investigation. Fifth, results in this current meta-analysis are generated from pre-to-post experiment, instead of between-group contrast. As a result, observed positive changes could be attributed to a variety of other factors that interact with the exercise interventions. For example, when people start an exercise program, they may change their diet and/or have more social interaction that are highly associated with improved cognitive function ^98–100^. Thus, future imaging studies should include active and/or passive control group (s) in order to draw a firm conclusion about the cognitive benefits of exercise intervention ^31^. Lastly, when using the well-recognized PEDro scale for risk of bias assessment, the majority of included studies scored 7 below (blinding of participants and instructor are unrealistic in exercise intervention, leading to 9 points in total). Thus, more well-designed randomized controlled trials should be further conducted on this topic.

## 5. Conclusions

The evidence for exercise effects on cognition is extensive but still growing. Combined with structural brain effects and behavioral data from previous studies, this article demonstrates that exercise-induced changes in functional brain activation in parietal regions (precuneus, superior and inferior parietal lobule, cingulate gyrus and posterior cingulate) and associated networks (frontoparietal network, dorsal attention network and default mode network) may neutrally mediate exercise-induced cognition enhancement. Furthermore, the present findings emphasize that the brain functional effects of exercise vary as a function of age and duration.

## Acknowledgement

The study is supported by no funding.

## Author Contributions

Q.Y. and L.Y.Z. planned the meta-analysis and formulated the hypotheses. Q.Y. and L.Y.Z performed the literature search and screening. Q.YS., L.Y.Z and Y.J.Z conducted the data extraction and coding, calculated the effect sizes and rated the quality. Q.Y., B.K.B. and L.Y.Z performed the statistical analyses. Q.Y. F.B, and L.Y.Z drafted the initial version of the manuscript. All authors were involved in revisions of the draft.

