## Supplementary data for "Cognitive Benefits of Exercise Interventions: An fMRI Activation Likelihood Estimation Meta-Analysis"

**Supplementary data 1.** Risk of bias assessment across 20 included studies

| Study | Score | Methodological Quality | PEDro Item Number | | | | | | | | | | |
|---|---|---|---|---|---|---|---|---|---|---|---|---|---|
| | | | 1 | 2 | 3 | 4 | 5 | 6 | 7 | 8 | 9 | 10 | 11 |
| Goldin et al., 2012a | 6 | Good | + | + | + | + | | | | + | | + | + |
| Goldin et al., 2012b | 6 | Good | + | + | + | + | | | | + | | + | + |
| Gourgouvelis et al., 2017 | 3 | Fair | + | | | | | | | + | | + | + |
| Schmitt et al., 2019 | 5 | Fair | + | + | | + | | | | + | | + | + |
| Boa Sorte Silva et al. 2020 | 6 | Good | + | + | | + | | | | + | + | + | + |
| Hsu et al., 2016 | 7 | Good | + | + | + | + | | | + | + | | + | + |
| Li et al. 2019 | 3 | Poor | + | | | | | | | + | | + | + |
| Martinsen et al. 2017 | 5 | Fair | + | + | | + | | | | + | | + | + |
| Nishiguchi et al. 2015 | 6 | Good | + | + | + | + | | | | + | | + | + |
| Smith et al. 2013 | 3 | Poor | + | | | | | | | + | | + | + |
| Krafft et al. 2014 | 5 | Fair | + | + | | + | | | | + | | + | + |
| Duchesne et al. 2016 | 6 | Good | + | + | + | + | | | | + | | + | + |
| Chen et al. 2016 | 4 | Fair | + | | | + | | | | + | | + | + |
| Pensel et al. 2018 | 4 | Fair | + | | | + | | | | + | | + | + |
| Liu-Ambrose et al. 2012 | 7 | Good | + | + | + | + | | | + | + | | + | + |
| Wagner et al. 2017 | 4 | Fair | + | | | + | | | | + | | + | + |
| Wu et al.2018 | 6 | Good | + | + | | + | | | | + | + | + | + |
| Wriessnegger et al. 2014 | 3 | Poor | + | | | | | | | + | | + | + |
| Metcalfe et al.2016 | 3 | Poor | + | | | | | | | + | | + | + |
| Baeck et al, 2012 | 4 | Fair | + | | | + | | | | + | | + | + |

Notes: (1) eligibility criteria; (2) random allocation; (3) concealed allocation; (4) similarity at baseline on key measures; (5) participant blinding; (6) instructor blinding; (7) assessor blinding; (8) more than 85% retention rate of at least one outcome; (9) intention-to-treat analysis; (10) between-group statistical comparison for at least one outcome; and (11) point estimates and measures of variability provided for at least one outcome. The studies were considered as excellent (9–10 points), good (6–8 points), fair (4–5 points), and poor (less than4 points) quality.

**Supplementary data 3**. Activation of subgroup analysis for health status at uncorrected level (uncorrected $p$ = 0.001, minimun volume = 200mm$^3$)

| Cluster | Activation Coordinate (MNI) | | | ALE value | Label (Nearest Gray Matter) |
|---|---|---|---|---|---|
| | x | y | z | | |
| | | | **Healthy Subgroup** | | |
| 1 | 24 | 6 | 0 | 0.014275 | Right Cerebrum. Sub-lobar. Lentiform Nucleus. Gray Matter. Putamen |
| 2 | 8 | -24 | 6 | 0.014021 | Right Cerebrum. Sub-lobar. Thalamus. Gray Matter. Pulvinar |
| 3 | 50 | 8 | 20 | 0.011956 | Right Cerebrum. Frontal Lobe. Inferior Frontal Gyrus. Gray Matter. Brodmann area 9 |
| 4 | -24 | -60 | -33 | 0.014508 | Left Cerebellum. Anterior Lobe. Gray Matter. |
| | | | **Unhealthy Subgroup** | | |
| 1 | 16 | -64 | 30 | 0.016207 | Right Cerebrum. Occipital Lobe. Precuneus. Gray Matter. Brodmann area 31 |
| 2 | -18 | -68 | 48 | 0.014174 | Left Cerebrum. Parietal Lobe. Precuneus. Gray Matter. Brodmann area 7 |
